# Transcripts enriched in codons that trigger the P-site tRNA–mediated mRNA decay possess stable mRNA

**DOI:** 10.1101/2025.04.10.648249

**Authors:** Rodolfo L. Carneiro, Fernando L. Palhano

## Abstract

Synonymous codon usage significantly influences mRNA stability in yeast by guiding mRNA decay during translation. The CCR4-NOT complex is central to this process, interacting with ribosomes when the A and E sites are unoccupied—a state that arises when a non-optimal codon with low tRNA availability is at the A site. This triggers recruitment of decay factors, reducing the stability of transcripts enriched in such codons. In humans, codon-mediated mRNA decay is less well understood. Recent research has identified a related but distinct mechanism called P-site tRNA-mediated decay (PTMD). Unlike yeast, human CCR4-NOT recruitment depends on specific arginine codons (CGG, CGA, or AGG) at the P site and slow decoding at the A site, allowing E-site vacancy and CNOT3-dependent binding. Through analysis of public datasets, we explored the characteristics of human transcripts enriched in PTMD codons. Interestingly, these codons are mostly found in transcripts with longer half-lives. This suggests that, rather than targeting already unstable mRNAs as in yeast, PTMD in humans selectively reduces the stability of otherwise long-lived transcripts, indicating a regulatory role distinct from the decay associated to codon usage.

## Introduction

Precise control of messenger RNA (mRNA) turnover is a key regulatory mechanism in numerous cellular processes. Some of these are inherent to mRNA function—for instance, transcripts with longer half-lives tend to accumulate in the cytoplasm and are translated into highly expressed proteins. Additionally, mRNA decay regulation has been linked to the immune response [1], responses to mRNA damage [2-5], and cell cycle progression and differentiation [6-8]. Various molecular machineries regulate mRNA decay, acting either globally or targeting specific motifs in transcript sequences [9].

The canonical pathway of mRNA degradation begins with deadenylation at the 3′ end by the Pan2–Pan3 and CCR4–NOT complexes [10-11], followed by decapping at the 5′ end by the Dcp1–Dcp2 complex. Once the mRNA ends are exposed, the exonuclease Xrn1 degrades the transcript in the 5′ to 3′ direction [12], while the Ski complex facilitates 3′ to 5′ degradation [13-14].

Although this canonical pathway is largely translation-independent, increasing attention has been given to translation-dependent regulation of mRNA stability, especially through synonymous codon usage. Despite the degeneracy of the genetic code, synonymous codons can influence mRNA half-life and protein expression in yeast [15]. Between the terms used to refer to codon usage, one key term is “codon optimality”, first defined in 2009 by Zhou, Weems and Wilke [17], as “the odds ratio of codon usage between highly and lowly expressed groups [of genes]”. Nevertheless, “optimality” has been used since at least 1985 to denote how well a codon-anticodon interaction occurs during translation, considering mainly tRNA availability [18]. Some codons are considered optimal because they match more abundant tRNAs, which leads to faster and more accurate translation. Others are non-optimal, being decoded by less abundant tRNAs, leading to a translation that is slower and more prone to errors [18]. Presnyak and colleagues demonstrated that synonymous codon optimality is a major determinant of mRNA stability. This relationship is mediated by the CCR4–NOT complex [16-19]. In yeast, the Not5 subunit of the CCR4– NOT complex interacts with the ribosomal E-site when both the A and E sites are unoccupied—a ribosome conformation that arises during slow decoding of non-optimal codons. This interaction promotes recruitment of the decapping factor Dhh1 and subsequent mRNA decay [19-20].

While the relation between codon optimality and mRNA decay is best characterized in yeast, it appears to be conserved across multiple species, including *Escherichia coli* [21], zebrafish [7-8], *Trypanosoma brucei* [22-23], *Drosophila melanogaster* [24], and humans [25-26].

In humans, however, the molecular mechanisms differ in key ways. A recent study revealed that CNOT3, the human homolog of yeast Not5, targets transcripts based on the structure of the tRNA occupying the ribosomal P-site [27]. Using ribosome footprint profiling in HEK293T cells, the authors showed that CNOT3 preferentially associates with ribosomes translating three specific arginine codons—CGA, CGG, and AGG—when both the A and E sites are vacant. This process, termed P-site tRNA-mediated decay (PTMD), requires both the presence of one of these codons in the P-site and slow decoding at the A-site to enable E-site vacancy and CCR4–NOT recruitment. Knockout of CNOT3 stabilizes numerous transcripts enriched in these codons, supporting its role in mRNA destabilization. PTMD is a codon-specific mRNA decay pathway [27]. The molecular mechanism that explain why CGA, CGG, and AGG codons trigger CCR4-NOT recruitment relays on the interactions between CNOT3, that is member of CCR4-NOT complex, and PTMD codons. Hydrogen bonding interactions with the D-arm of PTMD in the P-site promotes CNOT3 recruitment and mRNA degradation [27].

Despite these findings, it remains unclear whether transcripts enriched in PTMD codons differ in codon composition and mRNA stability compared to other transcripts. Moreover, the extent to which PTMD codons followed by poorly decoded codons influence transcript half-life is still not understood.

In this study, we investigate the genomic landscape of PTMD regulation by analyzing genes enriched in CGA, CGG, and AGG codons. Using datasets from Zhu and colleagues [27] and additional sources, we show that PTMD-target codons are primarily found in transcripts with long half-lives whereas unstable transcripts tend to lack PTMD codons. We also observed that potentially PTMD-target transcripts (rich in PTMD codons) have no preference for codons with low codon dwell time (fast translating) or high dwell time (slow translation) on A site followed the PTMD codon. Moreover, the presence of low or high codon dwell time on A site followed the PTMD codon have no effect into transcript stability. These findings suggest that PTMD has a mild impact in reducing half-lives of stable transcripts, rather than mediating decay of inherently unstable mRNAs.

## Results

### PTMD codons are enriched in stable transcripts

To investigate the possible roles of PTMD codons in the translation dynamics, particularly their effect on mRNA stability, we first quantified the frequency of the CGG, CGA and AGG (PTMD codons) within each transcript. Following the methodology used by Mendell’s group, we calculated a PTMD score for each transcript, weighting the CGG, CGA, and AGG codons by their enrichment in the P-site of CNOT3-associated ribosomes and normalizing by the total number of codons in the ORF [27]. This measure was defined as the transcript’s PTMD score. We then stratified the transcripts into two groups: the 1,000 transcripts with the highest PTMD scores (high PTMD group), and all remaining transcripts (low PTMD group) (Fig. 1A). We then checked if transcripts with high PTMD scores would have more stable mRNAs after CNOT3 knockout, if compared to genes with lower PTMD scores, using half-life data from Zhu and colleagues [27]. The high PTMD group presented a higher increase in mRNA half-life after CNOT3 knockout relative to the wild type cells (Fig. 1B) corroborating what was previously described [27]. Mendell’s group showed that the several mRNAs being destabilized by PTMD are mRNAs for highly expressed mitochondrial proteins [27]. To address if the translation pauses caused by signal peptides in transcripts addressing the mitochondrion would explain the result observed in Figure 1b, we performing the same analysis now discarding mitochondrial proteins. The difference between the two groups remains significant suggesting that PTMD codons, and not the signal peptides translation, are responsible for CNOT3 recruitment (Fig. S1). Interestingly, the high PTMD group also presented more stable mRNAs compared to the low-PTMD group, in both wild type cells (Fig. 1C) and CNOT3-KO cells (Fig. 1D).

**Figure 1.**
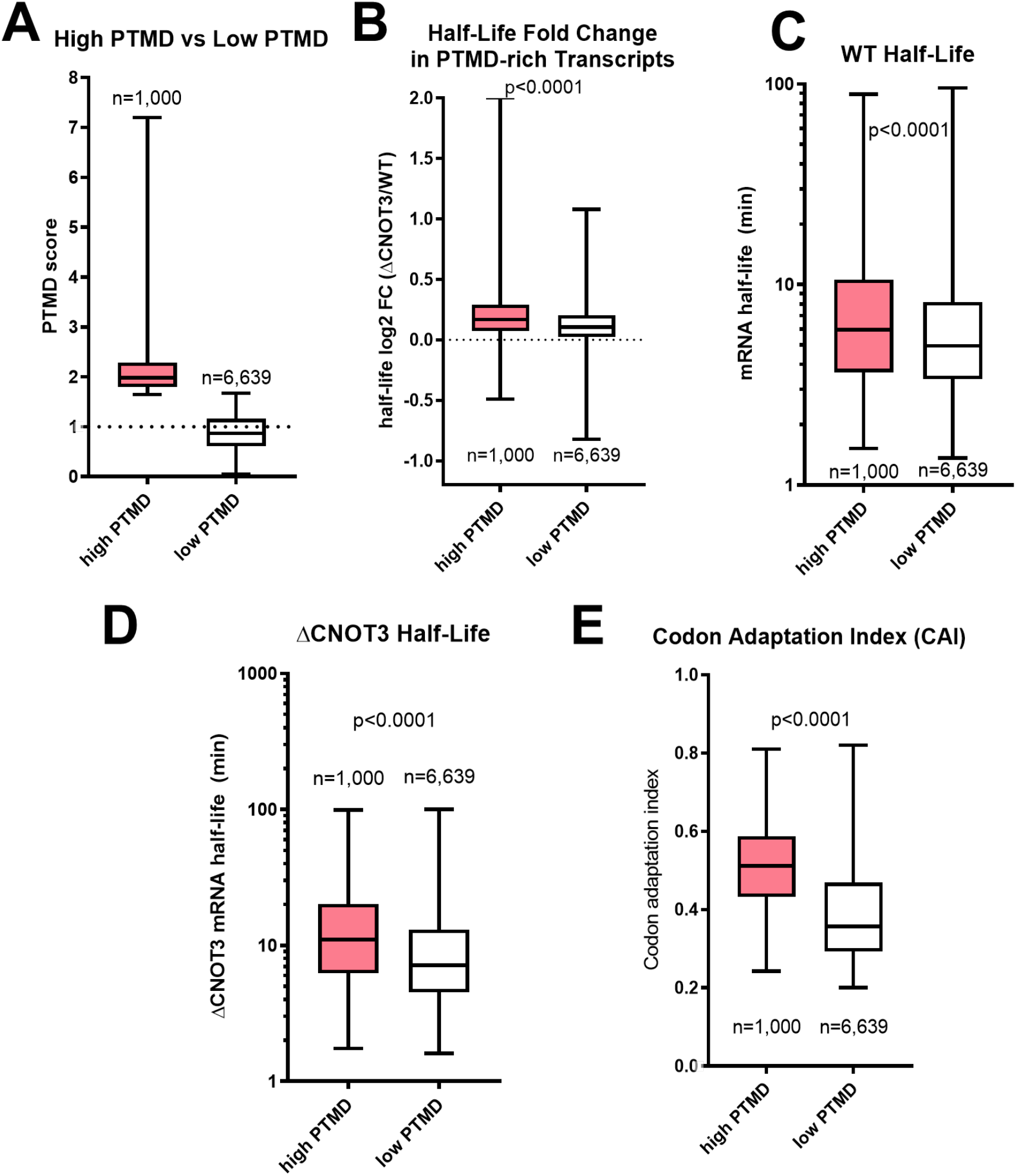
Comparison of PTMD score and transcripts rich in AGG, CGA, and CGG with other metrics. **(A)** Transcripts with high PTMD scores and other transcripts were split by their PTMD scores. **(B)** Log2 of the mRNA half-life fold change between CNOT3-knockout cells and wild-type cells. **(C)** mRNA half-life of transcripts with high PTMD score compared to mRNA half-life of the remaining transcripts in wild-type cells. **(D)** mRNA half-life of transcripts with high PTMD score compared to mRNA half-life of the remaining transcripts in CNOT3-knockout cells. **(E)** Codon Adaptation Index in transcripts with high PTMD score vs transcripts with low PTMD score. p-values given by Kolmogorov-Smirnov test.

Our data indicate that PTMD codons are more prevalent in stable mRNAs, instead of unstable ones, despite having a destabilizing effect. One possible explanation for this stability is the possibility of these transcripts being enriched in codons with stabilizing effect. Codon usage is known to influence mRNA stability, as codons that are more efficiently decoded tend to correlate positively with longer mRNA half-lives [26]. This property is often referred to as codon optimality. Several approaches can be used to estimate codon optimality depending on the context [17][26][28][29]. In this study, we used the codon adaptation index (CAI) as a proxy for codon optimality [28]. The CAI reflects how closely a transcript’s codon usage matches that of highly expressed human genes, thereby serving as an indicator of its potential translational efficiency [30]. Interestingly, transcripts in the high PTMD group also displayed higher CAI values than those in the low PTMD group (Fig. 1E).

In summary, we observed that transcripts enriched in PTMD codons are more stable and exhibit higher CAI values compared to the genome. Moreover, these transcripts show greater mRNA stabilization upon CNOT3 deletion, suggesting that they are preferential targets of the CCR4–NOT complex.

### Analyzing the dwell time and tRNA abundance of codons on the A-site following PTMD codons on the P-site

One of the key aspects of the PTMD mechanism is that CNOT3 binds to the ribosome only when both the E-site and A-site are vacant. Zhu and colleagues proposed that this occurs when the A-site codon has a long dwell time, giving enough time for the tRNA in the E-site to dissociate before a new aminoacyl-tRNA arrives at the A-site.

To explore whether this feature might have been evolutionarily selected as a mechanism for mRNA decay control, we investigated whether the dwell time of each codon correlates with how frequently it appears after the codons AGG, CGA, or CGG in the genome.

To do this, we developed a script to calculate the observed frequency of each codon pair in the genome and compare it to the expected frequency based on the individual usage of each codon. We refer to this metric as codon pair enrichment (shown on the y-axis of all panels in Figure 2).

**Figure 2.**
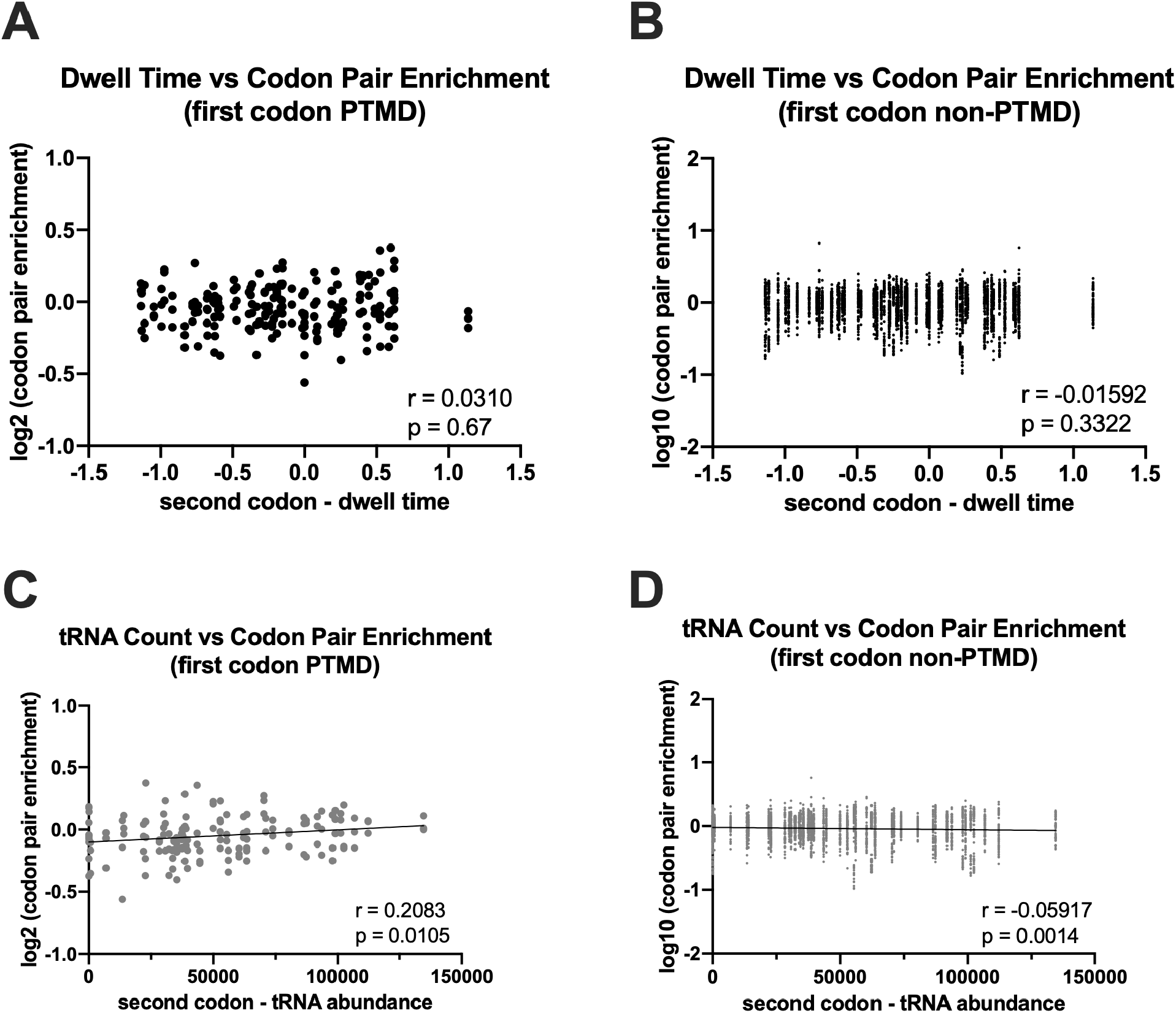
Correlations of PTMD occurrence, ribosome dwell time, and tRNA. **(A)** Correlation between codon pair occurrence in the genome ORFs, when the first codon of the pair is a PTMD codon, and the dwell time of the second codon in the pair. **(B)** Correlation between codon pair occurrence in the genome ORFs, when the first codon of the pair is a codon other than a PTMD one, and the dwell time of the second codon in the pair. **(C)** Correlation between codon pair enrichment in the genome ORFs, when the first codon of the pair is a PTMD codon, and the tRNA count for the anticodon correspondent to the second codon in the pair. **(D)** Correlation between codon pair occurrence in the genome ORFs, when the first codon of the pair is a codon other than a PTMD one, and the tRNA abundance for the anticodon correspondent to the second codon in the pair. Dwell time and tRNA abundance were obtained from [31] and [32] re-spectively.

In Figure 2A, codon pair enrichment was calculated for all 183 combinations (3 × 61) of codons, with one PTMD codon (AGG, CGA, or CGG) fixed in the first position. The x-axis represents the dwell time of the codon that follows the PTMD codon, as measured by ribosome profiling of HEK293T cells [31].

We found no correlation between the second codon’s dwell time and codon pair enrichment, regardless of whether the first codon was AGG, CGA, or CGG (Fig. 2A) or any other codon (Fig. 2B).

In contrast, when we used tRNA abundance [32] instead of dwell time, a statistically significant positive correlation (Pearson’s r > 0.2) was observed for codons following AGG, CGA, or CGG (Fig. 2C). For codons following non-PTMD codons, a weaker but significant negative correlation was found (Fig. 2D).

Finally, these effects appear to be specific to codons that follow AGG, CGA, or CGG, and not due to general proximity. When these PTMD codons were placed in the second position instead of the first, the correlations disappeared (Fig. S2).

A similar trend was observed when we used dwell time and tRNA abundance data from another study (Fig. S3) that analyzed hiPSC-derived neuronal cells [33]. We conclude that PTMD codons tend to be followed by codons with high tRNA abundance (Fig. 2C), although no bias was detected regarding the decoding time of these codons (Fig. 2A).

### CNOT3-associated mRNA decay is not dependent on ribosomal dwell time

To further clarify the impact of dwell time as well tRNA abundance on mRNA decay through PTMD, another script was used to search for each AGG, CGA and CGG in each transcript and calculate the average dwell time and average tRNA abundance of the codons following them. Then, we selected the 1,000 transcripts with the highest PTMD scores, the same ones as Fig. 1A, but we stratified them into two groups based on the median dwell time of the codons following AGG, CGA and CGG. For simplicity, these groups were called “high A-site DT” and “low A-site DT”, since the PTMD machinery is supposed to act when these codons with extreme dwell time values occupy the A-site of the ribosome. After stratifying our sample, we compared the mRNA half-life increase under CNOT3 knockout, in comparison to the wild type, between those two groups. Unexpectedly, the two groups showed no statistically significant difference (Fig. 3A).

**Figure 3.**
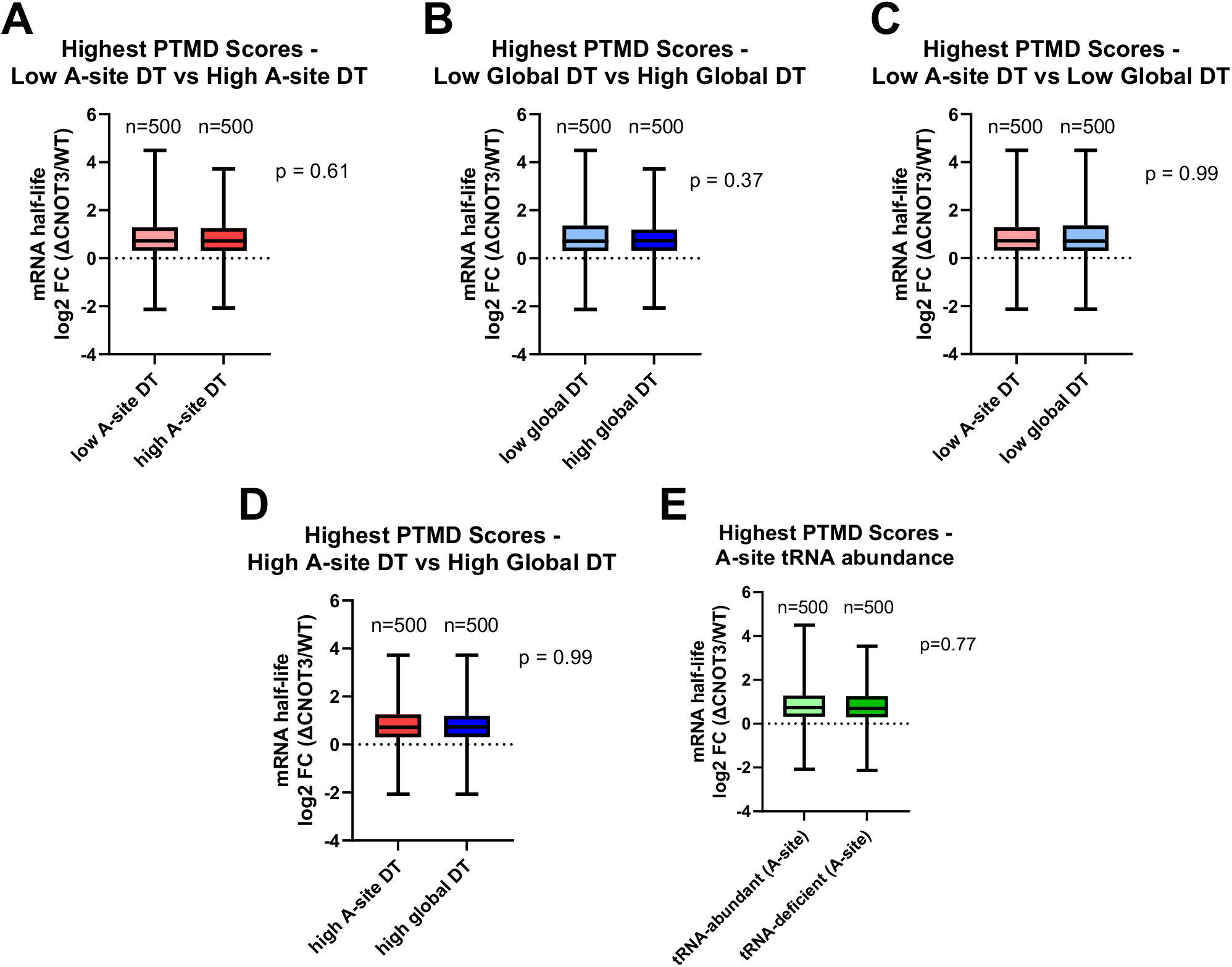
Dwell Time and tRNA availability of codons on the A-site have no impact on P-site tRNA Mediated Decay. (A) mRNA half-life fold change (ΔCNOT3/WT) of transcripts with high PTMD score comparing the lowest dwell time in the A-site after CGA, CGG or AGG vs transcripts with the highest dwell time in the A-site after CGA, CGG or AGG. (B) mRNA half-life fold change (ΔCNOT3/WT) of transcripts with high PTMD score, comparing the median dwell time for all codons throughout the transcript, instead of only the codons on the A-site following PTMD codons. (C) mRNA half-life fold change (ΔCNOT3/WT) of transcripts with high PTMD score, comparing transcripts with low dwell time on the A-sites following PTMD codons to the transcripts with low median dwell time for all the codons throughout the transcript. (D) mRNA half-life fold change (ΔCNOT3/WT) of transcripts with high PTMD score, comparing transcripts with high dwell time on the A-sites following PTMD codons to the transcripts with high median dwell time for all the codons throughout the transcript. (E) Same analysis as (A) but using tRNA abundance in place of dwell time. p-values given by Kolmogorov-Smirnov test. Dwell time and tRNA abundance were obtained from [31] and [32] respectively.

We next repeated the analysis, but stratifying the transcripts by the dwell time of every codon in their transcripts, instead of only the codons following AGG, CGA and CGG.

Again, the two groups showed no difference in the increase of mRNA half-life under knockout of CNOT3 (Fig. 3B). Subsequent comparisons between the groups with slow dwell time (Fig. 3C) or high dwell time (Fig. 3D), still demonstrated no statistically significant difference. Additionally, equivalent results were observed when substituting the dwell time by the tRNA abundance (Fig. 3E). Again, a similar scenario was observed when we used dwell time and tRNA abundance obtained from hiPSC neuronal cell type (Fig S4). In summary, features such as decoding time (Fig. 3A) and tRNA abundance (Fig. 3E) of codons following PTMD codons do not appear to influence the half-life of endogenous mRNAs after knockout of CNOT3.

### PTMD-rich and slow-decoding transcripts are not the most affected under CNOT3 knockout

After discussing how codon dwell time and tRNA abundance, in association with PTMD codons, affect a transcript’s half-life under CNOT3 knockout, we asked whether the transcripts most affected by CNOT3 knockout were the same we studied in the previous analysis. We then performed a cluster analysis comparing PTMD score, dwell time in the A-site after a PTMD codon, and half-life fold change after CNOT3 knockout, between 7,639 transcripts, grouping them by how similar they were regarding these variables. For simplicity, 7,639 transcripts were merged by k-means and clustered into 200 groups with similar values for these variables (Fig. 4). Interestingly, the codons with the most half-life change under CNOT3 knockout were clustered separately from those with the highest PTMD score and codon dwell time (Fig. 4), indicating that they don’t occur concomitantly, and other features than the PTMD machinery must be responsible for most of the effects of CNOT3 on mRNA half-life. This observation is consistent with the established role of CCR4-NOT as a broad regulator of mRNA stability [34-40].

**Figure 4.**
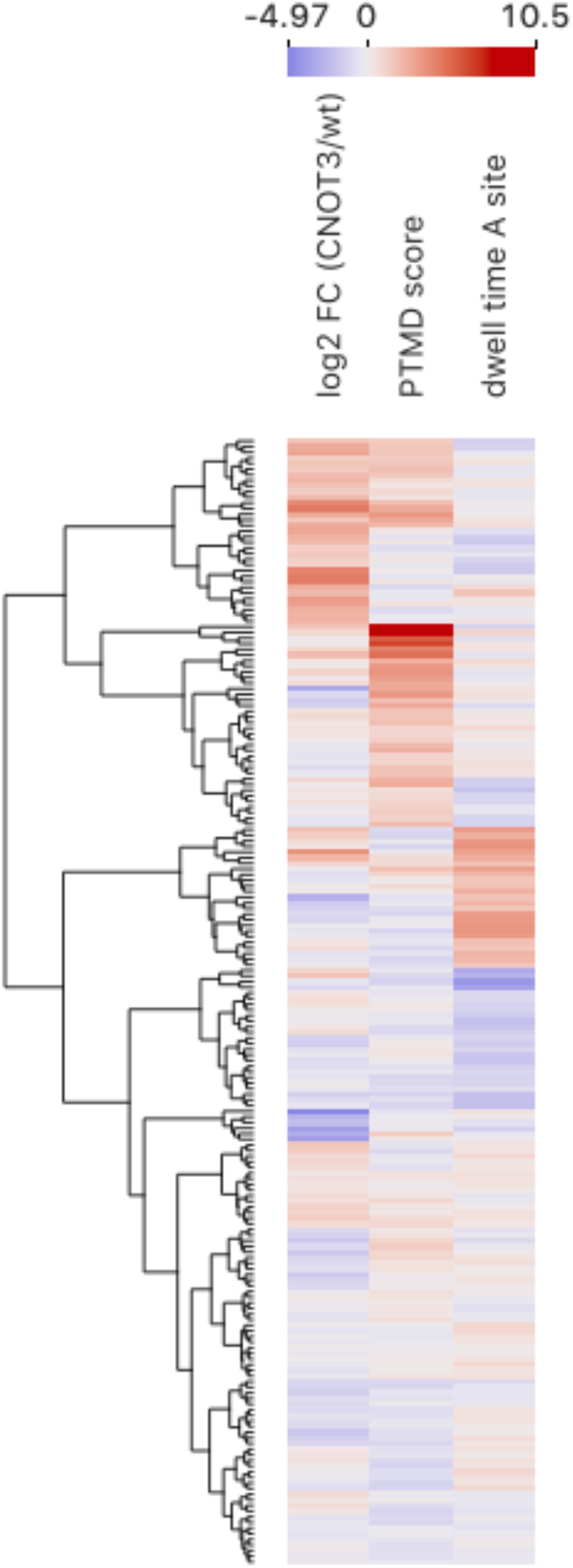
Cluster Analysis of PTMD score, mRNA half-life change after CNOT3 knockout, and A-site tAI or dwell time. Each 7,639 transcripts was grouped in one of 200 groups (k means), according to their similarity on the three metrics. Each line represents the average values of the transcripts in the group. The number of transcripts in each group may vary. The Values were normalized using Z-scores for each variable.

## Discussion

mRNA half-life correlation to codon optimality is conserved across various eukaryotic species, from yeast to humans, although being less prominent in the last one. The CCR4-NOT complex plays a key role in this correlation. Nevertheless, the complex seems to operate differently in humans than it does in yeast.

In yeast, the Not5 component of the CCR4-NOT complex targets slow decoding codons in the A-site. Codons with low tRNA availability are translated more slowly due to delayed codon-anticodon pairing, which promotes a ribosomal conformation with both the E-site and A-site vacant—this conformation is required for Not5 binding. Upon binding, Not5 recruits the mRNA decay machinery, resulting in shorter half-lives for transcripts enriched in slow-decoding codons.

Conversely, in humans, CNOT3—the homolog of Not5—preferentially targets specific codons in the P-site, particularly the AGG, CGA, and CGG codons of arginine. Although codon optimality is a major determinant of mRNA stability in yeast, its contribution to mRNA half-life in humans is less well understood. Bazzini and colleagues [7] demonstrated that codon identity is a major factor in human mRNA stability, independently of the poly-A tail. In contrast, Coller and colleagues highlighted the role of amino acid identity, in addition to codon bias, in mRNA regulation [26]. Alternatively, Agarwal and Kelley showed that other features contribute to the mRNA half-life as much as codon bias, such as ORF exon junction density and 3’ UTR length [41].

We observed that, despite the transcripts richer in AGG, CGA and CGG being more stable, this effect is highly independent from the CNOT3 effects through PTMD machinery, as proposed by Mendell [27]. Transcripts enriched in codon pairs where AGG, CGA, or CGG are immediately followed by a slow-decoding codon do not exhibit altered stability upon CNOT3 knockout when compared to transcripts with similar AGG, CGA, and CGG densities but followed by a rapid-decoding codon (Fig. 3A).

One possible explanation for this apparent discrepancy to Mendell’s results is the amount of PTMD codons required to recruit the CCR4-NOT complex. The minimal number of PTMD codons per transcript necessary to trigger a substantial mRNA degradation remains unknown. Mendell’s group used a reporter containing 42 tripeptides centered on a PTMD arginine codon to demonstrate accelerated CNOT3-mediated mRNA decay [27]. Interestingly, only 20 human genes closely resemble this reporter, containing at least 14 PTMD codons within a 42-codon window (data not shown). The extent to which PTMD contributes to global mRNA half-life regulation and the evolutionary basis for its specificity toward a subset of arginine codons remain to be elucidated.

## Methods

### Data sources

Coding sequences (CDS) and gene annotations for *Homo sapiens* GRCh38 were downloaded from Ensembl. Transcripts were filtered to include only those with lengths divisible by three and ending with a canonical stop codon. To avoid redundancy and overrepresentation of genes with multiple isoforms, only the longest transcript of each gene was retained. Of the 23,151 genes in the dataset, 17,324 (74.8%) had multiple annotated transcripts, with an average of 5.5 transcripts per gene. The selected longest transcripts had an average length of 564 codons and an average PTMD score of 0.193. In contrast, the discarded transcripts had an average length of 409 codons and an average PTMD score of 0.196.

Dwell time values correspond to the translation elongation times assigned to each of the 61 sense codons, as reported by Narula and colleagues [31], while the tRNA abundance values used here were obtained from Behrens and colleagues [32] (Figures 2 and 3). Alternatively, dwell time and tRNA abundance were obtained from [33] (Figures S3 and S4). The Codon Adaptation Index (CAI) values used in figure 1E were obtained from Subramanian and colleagues [42].

### Codon pair occurrence calculations

To quantify codon pair occurrence, we developed a script that enumerates the absolute frequency of each codon pair across the human transcriptome, considering all 3,904 possible combinations (61 × 64, excluding stop codons in the first position). Codon pairs were counted in-frame at any position within each transcript’s ORF.

The occurrence values used in Fig. 2 were obtained by dividing this absolute frequency by the pair expected frequency, given by the product of each codon frequency, as follows:

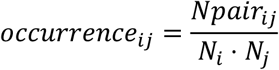

Where *Npair*_*ij*_ the frequency of the codon pair composed of codons *i* and *j*, and *N*_*i*_ and *N*_*j*_ representing the individual frequencies of codons *i* and *j*.

The data used for figure 2 is available in Supplemental Table 1.

### PTMD score

Zhu and colleagues [27] measured, through ribosome profiling, how often each codon was found occupying the ribosome P-site. This was done under CNOT3 immunoprecipitation and also in standard ribo-seq, and these values were used to calculate enrichment values for each codon. These values were called, here, as “weights” for the calculation of the PTMD score.

The PTMD score of a transcript was defined as the occurrence of AGG, CGA and CGG multiplied by the weight of the respective codon, and divided by the transcript length in codons, following the equation:

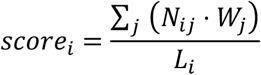

Where *score*_*i*_ is the PTMD score of the transcript *i, N*_*ij*_ is the number of times the codon *j* is found in the transcript *i, W*_*j*_ is the weight of the codon *j* as described above, and *L* is the length of transcript *i*, in codons.

### Search for PTMD codons and A-site codons following the PTMD codons

To identify transcripts containing PTMD target codons (AGG, CGA, or CGG) and the codons that appear immediately after them, we implemented a Python script that scans each coding sequence in our human transcriptome library. For every in-frame codon, the subsequent codon was considered to occupy the A-site relative to that P-site codon. In Figure 3, the dwell time and tRNA abundance values represent the median values of codons found in the A-site following each occurrence of AGG, CGA, or CGG. The median was chosen as the primary measure of central tendency due to the typically low frequency of PTMD codons per transcript, which can make the mean sensitive to outliers.

The data used for Fig. 1 A to D and Fig. 3 is available in Supplemental Table 2.

## Acknowledgments

We thank Joshua Mendell and Xiaoqiang Zhu for kindly sharing their data with us. We are also grateful to Janaina de Freitas Nascimento and Xiaoqiang Zhu for the critical reading of our manuscript.

## Funding

This work was supported by Conselho Nacional de Desenvolvimento Científico e Tecnológico (CNPq), Fundação de Amparo a Pesquisa do Estado do Rio de Janeiro (FAPERJ) and Coordenação de Aperfeiçoamento de Pessoal de Nível Superior (CAPES).

## Conflict of interest

The authors declare that they have no conflicts of interest with the contents of this article.

## Supplemental data

Supplemental data for this article can be accessed on the publisher’s website.

## Data availability

All data generated or analyzed during this study are included in this published article and its supplementary information files. The algorithms used during the current study are available from the corresponding author on reasonable request.

## Notes

### Competing Interest Statement

The authors have declared no competing interest.

### Summary of Updates

No updates were done. Mistakenly submitted to review.

