## Supplemental Figures for "Transcripts enriched in codons that trigger the P-site tRNA–mediated mRNA decay possess stable mRNA"

Rodolfo L. Carneiro and Fernando L. Palhano*
Programa de Biologia Estrutural, Instituto de Bioquímica Médica Leopoldo de Meis, Universidade Federal do Rio de Janeiro, Rio de Janeiro, RJ, 21941-902, Brazil.

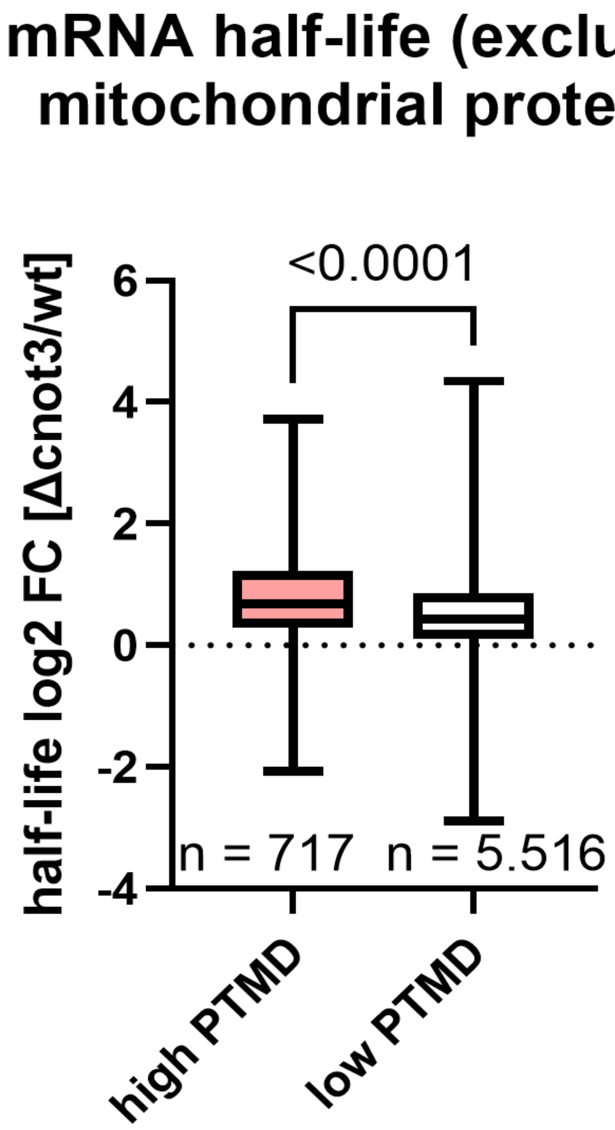


**Figure S1. Half-life fold change under CNOT3 knockout over the wild-type half-life, in log2.** The two groups were stratified the same way as figure 1, but excluding any transcript addressed as mitochondrial by its gene ontology.

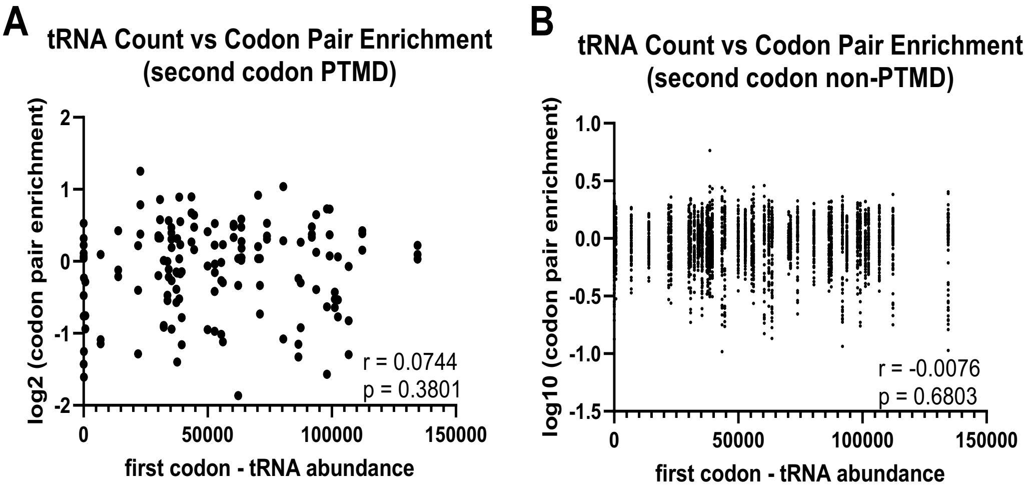


**Figure S2. Pearson's correlation coefficient between codon pair enrichment and tRNA abundance of the first codon of the pair (A)** when the second codon of the pair is a PTMD codon, or **(B)** when the second codon of the pair is not a PTMD codon.

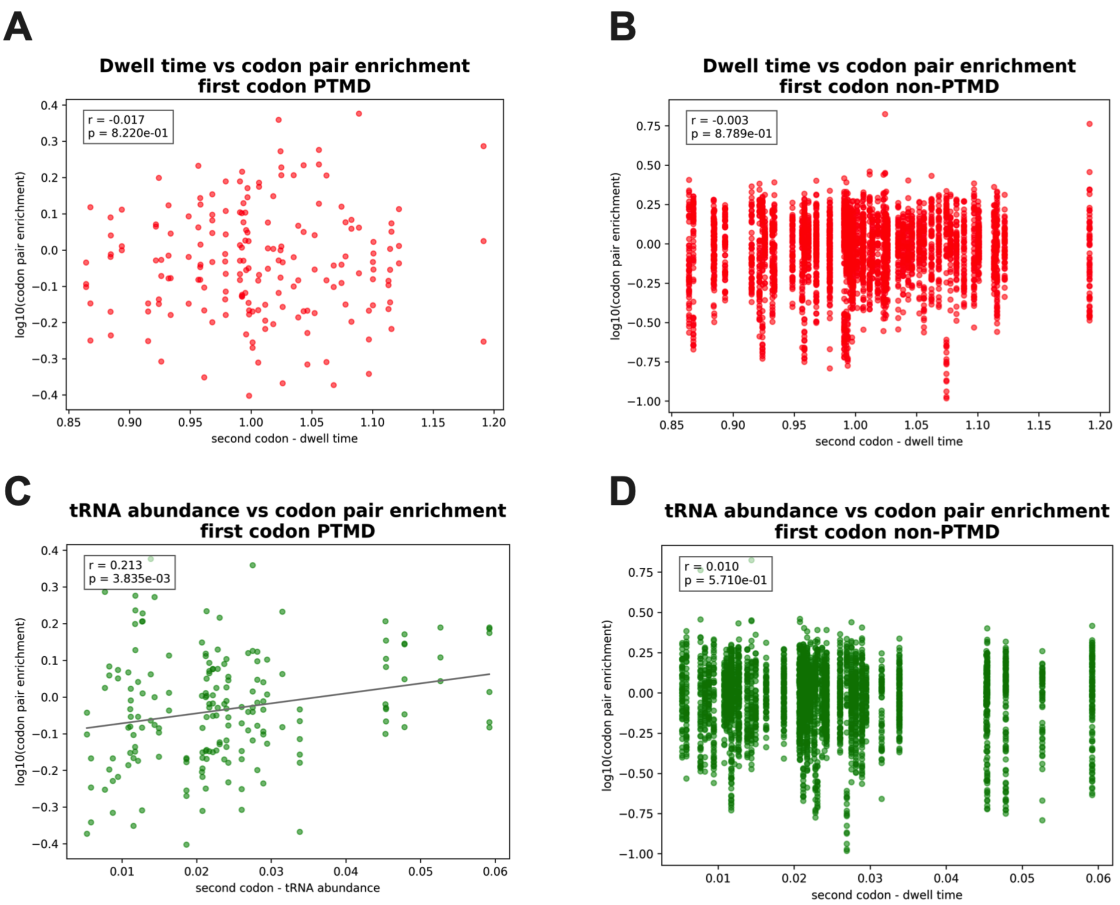


**Figure S3. Correlations of PTMD occurrence, ribosome dwell time, and tRNA. (A)** Correlation between codon pair occurrence in the genome ORFs, when the first codon of the pair is a PTMD codon, and the dwell time of the second codon in the pair. **(B)** Correlation between codon pair occurrence in the genome ORFs, when the first codon of the pair is a codon other than a PTMD one, and the dwell time of the second codon in the pair. **(C)** Correlation between codon pair enrichment in the genome ORFs, when the first codon of the pair is a PTMD codon, and the tRNA count for the anticodon correspondent to the second codon in the pair. **(D)** Correlation between codon pair occurrence in the genome ORFs, when the first codon of the pair is a codon other than a PTMD one, and the tRNA abundance for the anticodon correspondent to the second codon in the pair. Dwell time and tRNA abundance were obtained from [33].

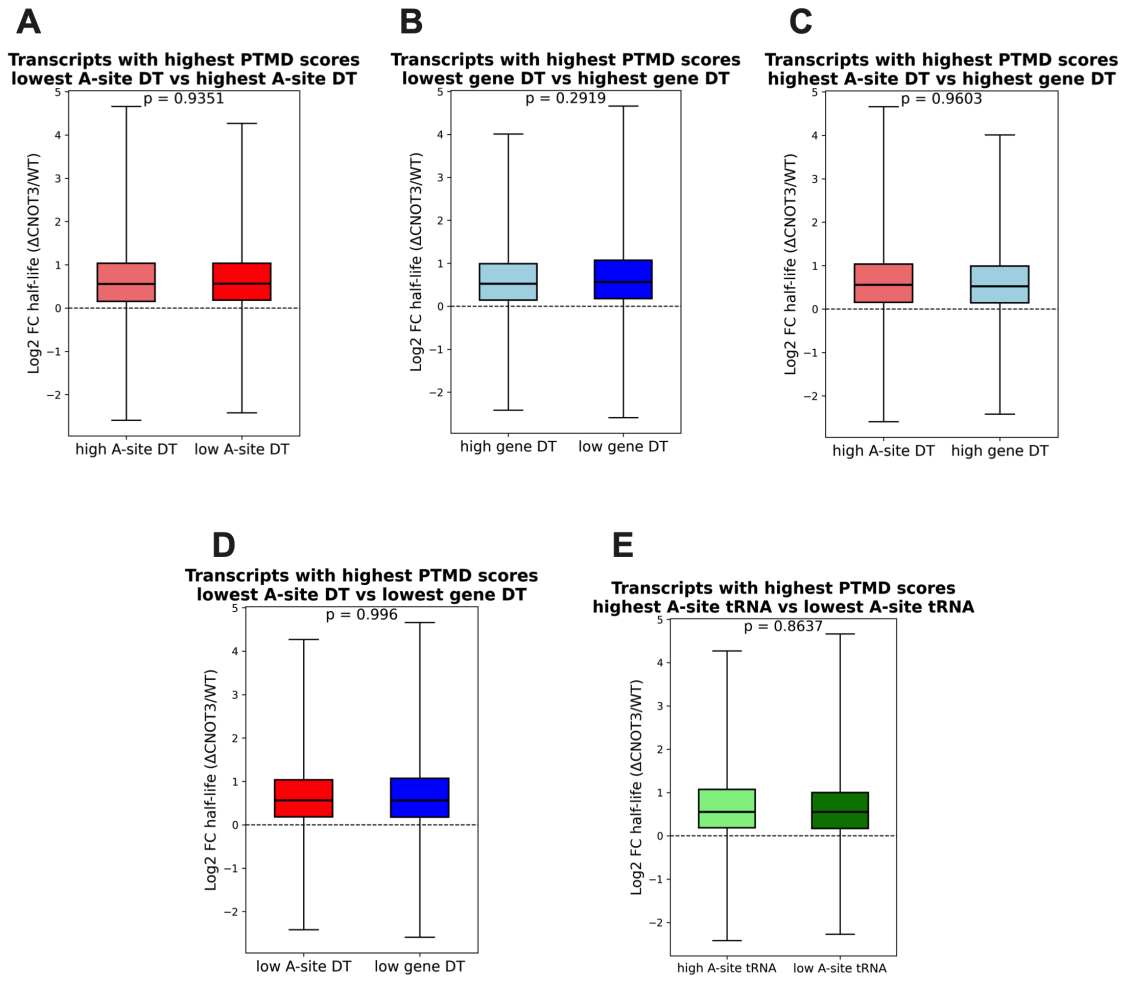


**Figure S4. Dwell Time and tRNA availability of codons on the A-site have no impact on P-site tRNA Mediated Decay.** (A) mRNA half-life fold change (ΔCNOT3/WT) of transcripts with high PTMD score comparing the lowest dwell time in the A-site after CGA, CGG or AGG vs transcripts with the highest dwell time in the A-site after CGA, CGG or AGG. (B) mRNA half-life fold change (ΔCNOT3/WT) of transcripts with high PTMD score, comparing the median dwell time for all codons throughout the transcript, instead of only the codons on the A-site following PTMD codons. (C) mRNA half-life fold change (ΔCNOT3/WT) of transcripts with high PTMD score, comparing transcripts with low dwell time on the A-sites following PTMD codons to the transcripts with low median dwell time for all the codons throughout the transcript. (D) mRNA half-life fold change (ΔCNOT3/WT) of transcripts with high PTMD score, comparing transcripts with high dwell time on the A-sites following PTMD codons to the transcripts with high median dwell time for all the codons throughout the transcript. (E) Same analysis as (A) but using tRNA abundance in place of dwell time. p-values given by Kolmogorov-Smirnov test. Dwell time and tRNA abundance were obtained from [33].
